# Persistent TOP1 cleavage complexes in drug-tolerant cells drive adaptive resistance to EGFR-targeted therapies in lung cancer

**DOI:** 10.64898/2026.06.02.729207

**Authors:** Mathéa Geraud, Remi Gence, Anne Casanova, Estelle Taranchon-Clermont, Simona Salimbeni, Nesibe Ozsu, Maina Vienne, Célia Delahaye, Elodie Borrull, Amélie Lusque, Mathilde Morisseau, Thomas Filleron, Delphine Pagan, Camille Taha, Agnese Cristini, Julien Mazières, Olivier Calvayrac, Gilles Favre, Anne Pradines, Olivier Sordet

## Abstract

Resistance to targeted cancer therapies often arises from drug-tolerant cells (DTCs), which survive treatment by entering a non-proliferative state. Over time, DTCs can acquire mutations that contribute to cell reproliferation, but how non-proliferating DTCs accumulate such mutations remains unclear. Here, we show that EGFR inhibition in EGFR-mutated lung cancer transiently downregulates tyrosyl-DNA phosphodiesterase 1 (TDP1), a repair enzyme that resolves abortive topoisomerase I cleavage complexes (TOP1ccs). In DTCs, elevated reactive oxygen species promote TOP1cc trapping, while TDP1 downregulation impairs their repair, driving TOP1cc accumulation, resistance mutation acquisition, and cell reproliferation. We further find that TDP1 expression is absent in approximately 25% of EGFR-mutated lung cancers. In TDP1-deficient cells, combining EGFR inhibition with a sublethal concentration of topotecan, which further increases TOP1ccs, abolishes cell reproliferation. Together, these findings establish persistent TOP1cc accumulation as a driver of therapy-induced mutagenesis linking drug tolerance to adaptive resistance, and reveal TDP1 loss as a targetable vulnerability in EGFR-mutated lung cancers.

**Teaser:** A drug-tolerant state that fuels mutagenesis and resistance also creates a transient vulnerability.

## INTRODUCTION

Targeted therapies exploit tumor-specific genetic alterations on which cancer cell survival depends. Targeting EGFR alterations in non-small cell lung cancer (NSCLC) represents a paradigm of targeted therapy, providing substantial benefit over standard chemotherapy (*1*). However, despite initial and sometimes prolonged responses, resistance inevitably emerges, leading to tumor recurrence (*2-6*). Acquired resistance can arise from a small subset of tumor cells that survive treatment by entering a quiescent or dormant state (*7, 8*). These drug-tolerant cells (DTCs) emerge rapidly upon drug exposure and constitute a reversible, non-genetic adaptive state that enables survival under therapeutic pressure (*2-6*). DTCs display marked phenotypic plasticity, including transcriptional reprogramming (*9-11*) and metabolic adaptations (*10*). Over time, however, they can transition to stable resistance through diverse mechanisms, including the acquisition of de novo genetic alterations that restore proliferative capacity under treatment (*12*). Studies in NSCLC and other cancers, including colorectal carcinoma, suggest that DTCs may enter a transiently mutagenic state through downregulation of high-fidelity DNA repair pathways, such as homologous recombination (HR) (*13*) and base excision repair (BER) (*14*), coupled with upregulation of error-prone processes, including translesion DNA synthesis (*13*) and cytidine deaminase activity (*15-17*). Yet, how non-proliferating DTCs acquire secondary resistance mutations, and how this mutagenic state is mechanistically linked to the drug-tolerant program, remain poorly understood.

DNA single-strand breaks (SSBs) are the most frequent endogenous DNA lesions, and defective SSB repair is a major source of genome instability (*18, 19*). In non-proliferative but transcriptionally active cells, an important source of SSBs arises from DNA topoisomerase I (TOP1), which relieves transcription-associated torsional stress by generating transient TOP1-DNA cleavage complexes (TOP1ccs), in which TOP1 is covalently linked to the 3′ end of a SSB (*20, 21*). Although TOP1ccs are normally short-lived and rapidly reversed upon DNA relaxation, frequent physiological and pathological DNA alterations can impair TOP1 religation, leading to abortive TOP1ccs in which TOP1 remains covalently trapped on DNA and the SSB persists (*21, 22*). These trapped TOP1ccs are intrinsically mutagenic and therefore require efficient repair to preserve genome stability (*22-24*). Their resolution primarily depends on proteasome-mediated processing of TOP1, followed by excision of the residual TOP1 peptide by tyrosyl-DNA phosphodiesterase 1 (TDP1), which generates TOP1-free SSB intermediates that are subsequently sealed by the XRCC1-PNKP-LIG3 repair machinery (*25-27*). Consistent with a critical role in non-dividing cells, inactivating mutations in *TDP1* cause the neurodegenerative syndrome spinocerebellar ataxia with axonal neuropathy 1 (SCAN1), characterized by progressive neuronal loss (*28-30*). However, whether defective TOP1cc repair contributes to mutagenesis and resistance evolution in DTCs remains unknown.

Here, we identify impaired TOP1cc repair as a mechanism that links adaptive drug tolerance to mutagenesis under EGFR-targeted therapy. We show that transient downregulation of TDP1, together with elevated oxidative stress, promotes the accumulation of abortive TOP1ccs in DTCs, thereby fostering mutation acquisition and re-entry into proliferation under treatment. Moreover, we uncover TDP1 deficiency as a therapeutic vulnerability, as pharmacologic induction of TOP1ccs in combination with EGFR inhibition suppresses the emergence and re-proliferation of resistant cells.

## RESULTS

### Erlotinib transiently downregulates the TDP1 repair pathway

To investigate the role of TDP1 in secondary resistance to targeted therapy, we used the EGFR-mutated NSCLC cell line PC9 treated with the first-generation EGFR-tyrosine kinase inhibitor (EGFR-TKI) erlotinib. Although osimertinib is currently the standard-of-care EGFR-TKI in clinical settings, first-generation EGFR-TKI models are widely used and well characterized to study acquired resistance, as they recapitulate key resistance mechanisms observed across EGFR inhibitors (*2, 7-9, 11, 12, 15*). PC9 cells were subcloned (*9*) to reduce heterogeneity and minimize the presence of pre-existing resistant cells (*8, 31*). High-content microscopy of individual PC9 cells treated with erlotinib, revealed three sequential phases (Fig. 1A), consistent with previous reports (*7, 8, 11*): an initial rapid decrease in cell number reflecting elimination of the majority of cells, followed by a plateau corresponding to a small population of quiescent/dormant DTCs, and subsequently, the re-emergence of proliferating drug-tolerant expanded clones (DTECs) (Fig. 1A).

**Fig. 1.**
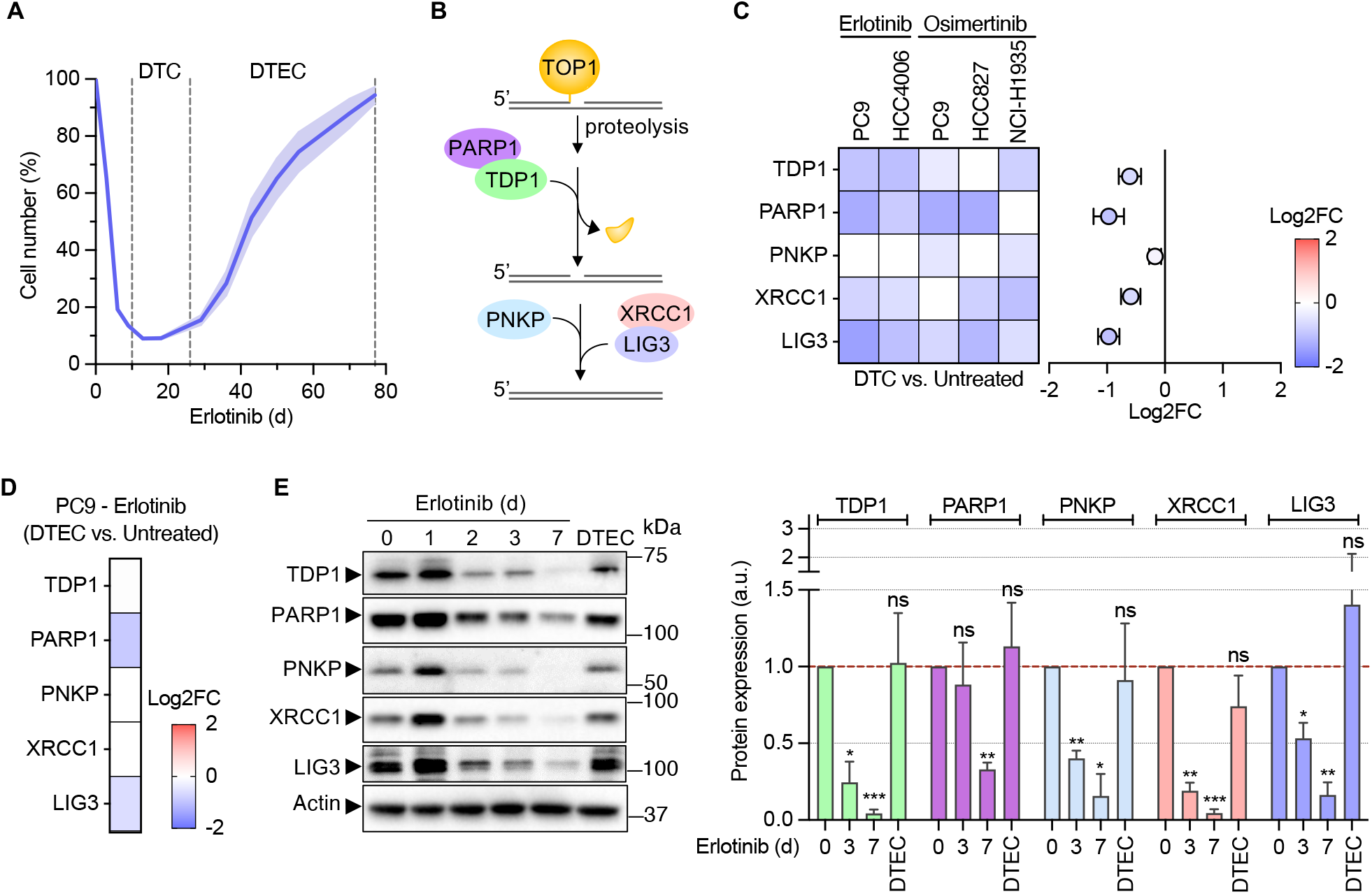
EGFR-TKI treatment transiently decreases the expression of TDP1 repair complex components. (**A**) Time course of PC9 cell number in response to erlotinib (1 µM), monitored by phase-contrast imaging and normalized to untreated cells (100%). A representative experiment out of seven independent experiments is shown (mean ± SEM; n = 6 technical replicates). The bold line represents the mean, and the shaded area the SEM. DTC and DTEC denote cellular stages based on the curve dynamics. (**B**) Schematic of TOP1cc removal by the TDP1 repair pathway. (**C)** Differential expression of TDP1 complex genes in PC9, HCC4006, HCC827, and NCI-H1975 cells treated with erlotinib or osimertinib. Heatmap shows log_2_ fold changes (log2FC) of transcript levels in DTC relative to untreated cells. White boxes indicate non-significant changes (adjusted *P*-value ≥ 0.05). Right panel shows mean log2FC ± SEM for each transcript across the five cell line/treatment combinations shown in the left panel (n = 5). (**D)** Heatmap as in (C) but showing transcript levels in DTEC (several weeks of continuous culture with 1 µM erlotinib following relapse) relative to untreated PC9 cells. (**E**) Western blot of proteins of the TDP1 complex in PC9 cells treated with erlotinib (1 µM) for the indicated times and in DTEC. Actin: loading control. Left: representative blots. Right: quantification of the indicated proteins normalized to actin (mean ± SEM; n = 3). ns, not significant, \**P* < 0.05, \*\**P* < 0.01, **\*\*\****P* < 0.001 (one-sample t-test vs. 1).

In erlotinib-treated PC9 cells, we performed RNA-seq that revealed marked downregulation of multiple components of the TDP1-mediated TOP1cc repair pathway, including TDP1, PARP1, PNKP, XRCC1, and LIG3 after 7 days of treatment (Fig. 1B,C). This downregulation coincided with G0/G1 arrest (Fig. S1A) and entry into the DTC state (Fig. 1A). Analysis of publicly available databases (*9, 32*) showed a similar reduction in RNA levels in several EGFR-mutated NSCLC cell lines treated with erlotinib (HCC4006) or osimertinib (PC9, HCC827, NCI-H1975) at the DTC state (Fig. 1C), indicating that downregulation of the TDP1 pathway is a conserved feature of EGFR-TKI-induced DTC state. This decrease was transient, as transcript levels returned fully or partially to baseline in the DTECs from erlotinib-treated PC9 cells (Fig. 1D). Consistently, protein levels of all components of the TDP1 pathway were reduced over the first 7 days of treatment and restored in DTECs (Fig. 1E), confirming the transient nature of this pathway downregulation. WI38-hTERT cells induced into quiescence displayed a reduction, albeit weaker, in the expression of the same repair components compared with proliferative cells (Fig. S1B), suggesting that cell cycle exit could contribute to the downregulation of the TDP1-mediated repair pathway observed in DTCs. Together, these results indicate that TDP1 pathway downregulation is a transient feature of the EGFR-TKI-induced DTCs and is restored upon acquisition of resistance.

### Erlotinib-induced TDP1 downregulation promotes TOP1cc accumulation

The TDP1 pathway removes trapped TOP1ccs (*25, 26*), raising the question of whether its downregulation upon erlotinib treatment leads to TOP1cc accumulation. TOP1ccs were detected as discrete foci by immunofluorescence microscopy using an anti-TOP1cc antibody (*30, 33*). In PC9 cells, erlotinib treatment progressively increased the number of TOP1cc foci over 7 days (Fig. 2A), and a similar trend was observed in HCC4006 cells (Fig. S2A). In PC9 cells, this accumulation paralleled the progressive downregulation of TDP1 pathway components (Fig. 1C-E) as cells entered the DTC state (Fig. 1A, Fig. S1A). In DTECs, where TDP1 pathway components were re-expressed (Fig. 1D,E), TOP1cc foci returned to baseline (Fig. 2A). Notably, TOP1 expression transiently decreased during treatment (Fig. S2B), indicating that TOP1cc accumulation occurs despite lower enzyme abundance.

**Fig. 2.**
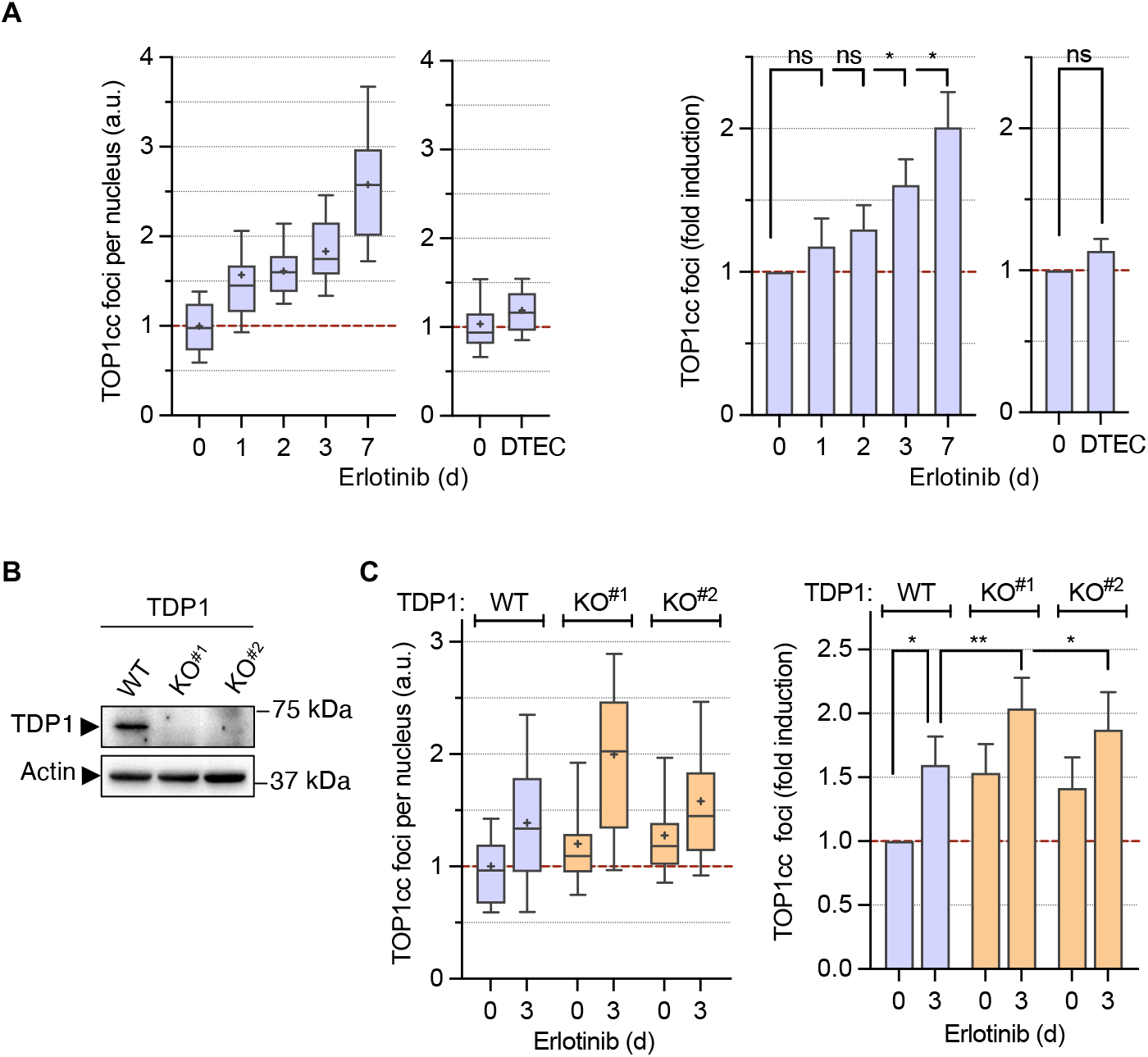
Erlotinib treatment transiently increases TOP1ccs. (**A**) PC9 cells were treated with erlotinib (1 µM) for the indicated times and stained for TOP1ccs and DAPI (DNA). DTEC corresponds to several weeks of continuous culture with erlotinib (1 µM) following relapse. Number of TOP1cc foci per nucleus normalized to untreated cells in a representative experiment (left) and in three to four independent experiments (right: mean ± SEM; n = 3-4). ns, not significant, \**P* < 0.05 (two-tailed paired t-test on non-normalized log2-transformed values). (**B**) Western blot of TDP1 in TDP1 WT and KO PC9 cells. Actin: loading control. (**C**) Same experiment as in (A) in TDP1 WT and KO PC9 cells treated with erlotinib (1 µM) for 3 days (mean ± SEM; n = 5). \**P* < 0.05, \*\**P* < 0.01 (two-tailed paired t-test on non-normalized log2-transformed values).

To directly test the impact of TDP1 downregulation on adaptive resistance, we generated TDP1 knockout (KO) PC9 cells using CRISPR-Cas9 (Fig. 2B). TDP1 depletion did not affect neither cell growth (Fig. S2C), nor the expression of TOP1 (Fig. S2D), and other TDP1 pathway components (Fig. S2E). Consistent with the established role of TDP1 in TOP1cc repair (*25, 26, 30*), TDP1 KO cells showed further TOP1cc accumulation upon erlotinib treatment compared to wild-type (WT) cells (Fig. 2C). Thus, transient TDP1 pathway downregulation in response to erlotinib promotes TOP1cc accumulation, likely reflecting impaired repair capacity.

### Erlotinib-induced TDP1 downregulation accelerates EGFR T790M acquisition and DTC outgrowth

Trapped TOP1ccs are a well-established source of mutations and genomic instability (*22-24*). In PC9 cells and in patients, acquired resistance to erlotinib frequently arises through secondary mutations, with the EGFR T790M substitution accounting for approximately half of resistance-associated events (*12, 34*). This well-characterized and recurrent mutation in this model provides a robust and quantifiable readout of mutation acquisition. We therefore asked whether TDP1 downregulation-mediated TOP1cc accumulation promotes the emergence of EGFR T790M in DTCs. To test this, we compared TDP1 WT with TDP1 KO cells to further reduce TDP1 expression and thereby increase TOP1cc levels.

EGFR T790M was quantified by digital droplet PCR (ddPCR), a highly sensitive method for mutation detection (*35*), after 10-14 days of erlotinib treatment. At this time, WT and TDP1 KO cells remained in a comparable DTC state, as indicated by robust G0/G1 arrest (Fig. S1A), and similar cell numbers (Fig. S3A) and sensitivity to erlotinib (Fig. S3B). EGFR T790M remained below the ddPCR detection limit in untreated WT and TDP1 KO cells, but became detectable following erlotinib treatment (Fig. 3A), consistent with previous reports indicating that T790M is acquired during drug exposure rather than pre-existing in this model (*7, 8, 12*). Across independent experiments, TDP1 KO cells showed a greater increase in EGFR T790M fractional abundance than WT cells, with a more pronounced and reproducible effect in TDP1 KO^#2^ (Fig. 3A). Although the magnitude of T790M emergence varied across experiments, consistent with the stochastic nature of resistance mutation acquisition, these data indicate that TDP1 loss accelerates the acquisition of the EGFR T790M resistance mutation during the DTC state.

**Fig. 3.**
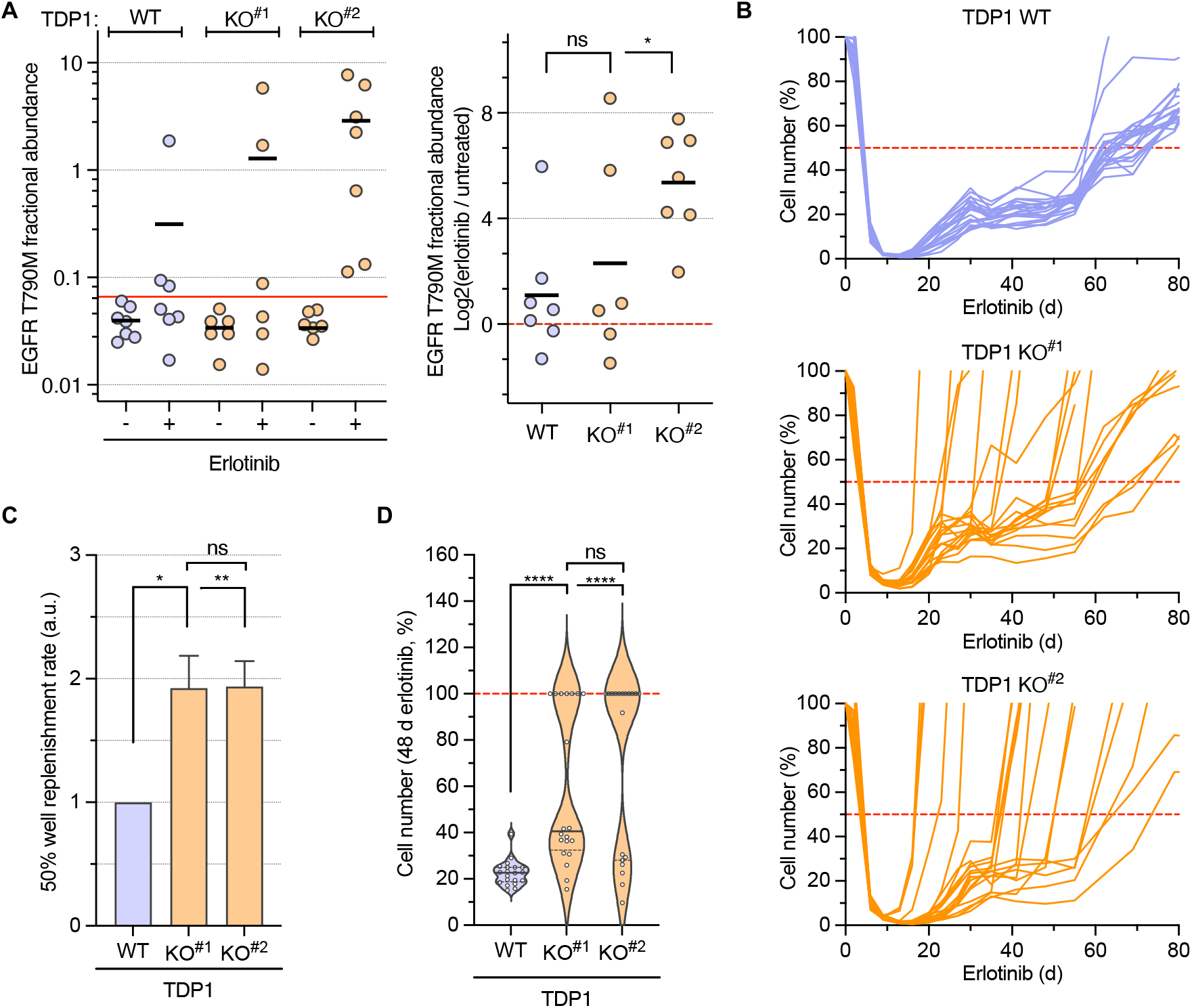
TDP1 depletion accelerates the acquisition of the EGFR T790M mutation and the resumption of cell proliferation in response to erlotinib. (**A**) TDP1 WT and KO PC9 cells were treated with erlotinib (1 μM) for 10-14 days. Left: ddPCR quantification of EGFR T790M fractional abundance. Each dot represents the mean of technical triplicates from one independent experiment (n = 6-7). Red line denotes the limit of detection (LOD). Right: log2 fold-change (log2FC) of EGFR T790M fractional abundance upon erlotinib treatment relative to matched untreated controls per experiment. Each dot represents one independent experiment. ns, not significant, \**P* < 0.05 (two-tailed Wilcoxon matched-pairs signed-rank test). (**B**) Time course of TDP1 WT and KO PC9 cell number upon erlotinib treatment, monitored by phase-contrast imaging and normalized to untreated cells (100%). Each line represents one technical replicate (n = 20). A representative experiment out of three independent experiments is shown. (**C**) Quantification of the three independent experiments shown in (B). Data are normalized to TDP1 WT cells and expressed as 1/median time to reach 50% well replenishment (mean ± SEM; n = 3). ns, not significant, \**P* < 0.05, \*\**P* < 0.01 (two-tailed paired t-test on non-normalized log2-transformed median times). (**D**) Violin plot representation of the same representative experiment shown in (B) at day 48 of erlotinib treatment. Each dot represents one technical replicate. ns, not significant, \*\*\*\**P* < 0.0001 (two-tailed unpaired t-test).

Because the EGFR T790M mutation confers resistance to erlotinib and enables DTC outgrowth (*7, 8, 12*), we next asked whether its earlier emergence in TDP1 KO cells translated into earlier reproliferation. Consistent with this possibility, TDP1 KO cells resumed proliferation earlier than WT cells during prolonged erlotinib exposure, as reflected by faster repopulation of the culture (Fig. 3B). The median time to well replenishment was approximately twofold shorter in TDP1 KO cells than in WT cells (Fig. 3C). Notably, accelerated reproliferation was observed in only approximately half of the TDP1 KO wells analyzed (Fig. 3D), consistent with heterogeneous resistance emergence across independent wells. Earlier reproliferation of TDP1 KO cells was not due to faster growth of resistant cells, as DTECs derived from TDP1 KO and WT cells showed comparable proliferation rates (Fig. S3C), supporting the interpretation that TDP1 loss accelerates the acquisition of resistance during the DTC state. Together, these findings support a model in which erlotinib-induced TDP1 downregulation promotes TOP1cc accumulation, thereby accelerating EGFR T790M emergence and earlier DTC outgrowth.

### ROS induce TOP1cc accumulation and DTC outgrowth during erlotinib treatment

TOP1ccs can be trapped by reactive oxygen species (ROS)-induced oxidative DNA lesions (*36-38*), and the EGFR-TKIs gefitinib and osimertinib have been reported to increase intracellular ROS (*10, 39*), We therefore asked whether ROS contribute to erlotinib-induced TOP1cc accumulation and subsequent DTC outgrowth.

Analysis of erlotinib-treated PC9 cells using the luminescent ROS reporter ROS-Glo revealed a marked increase in ROS after 3 days of treatment (Fig. 4A). Consistent with increased oxidative stress, erlotinib also induced the accumulation of 8-oxoguanine (8-oxoG) at this time point (Fig. 4B). In contrast, no ROS increase was detected in DTECs (Fig. 4C), paralleling the transient accumulation of TOP1ccs observed during erlotinib treatment (Fig. 2). To directly test whether ROS induce TOP1cc accumulation, cells were treated with the ROS scavenger N-acetylcysteine (NAC). NAC completely suppressed both erlotinib-induced ROS (Fig. 4A) and TOP1cc accumulation (Fig. 4D). Similar results were obtained in TDP1 KO PC9 cells (Fig. S4A,B), supporting a major role of ROS in erlotinib-induced TOP1cc accumulation. We next asked whether ROS also contribute functionally to DTC outgrowth during prolonged erlotinib treatment. NAC significantly reduced reproliferation of TDP1 WT cells and completely suppressed the enhanced reproliferation observed in TDP1 KO cells (Fig. 4E).

**Fig. 4.**
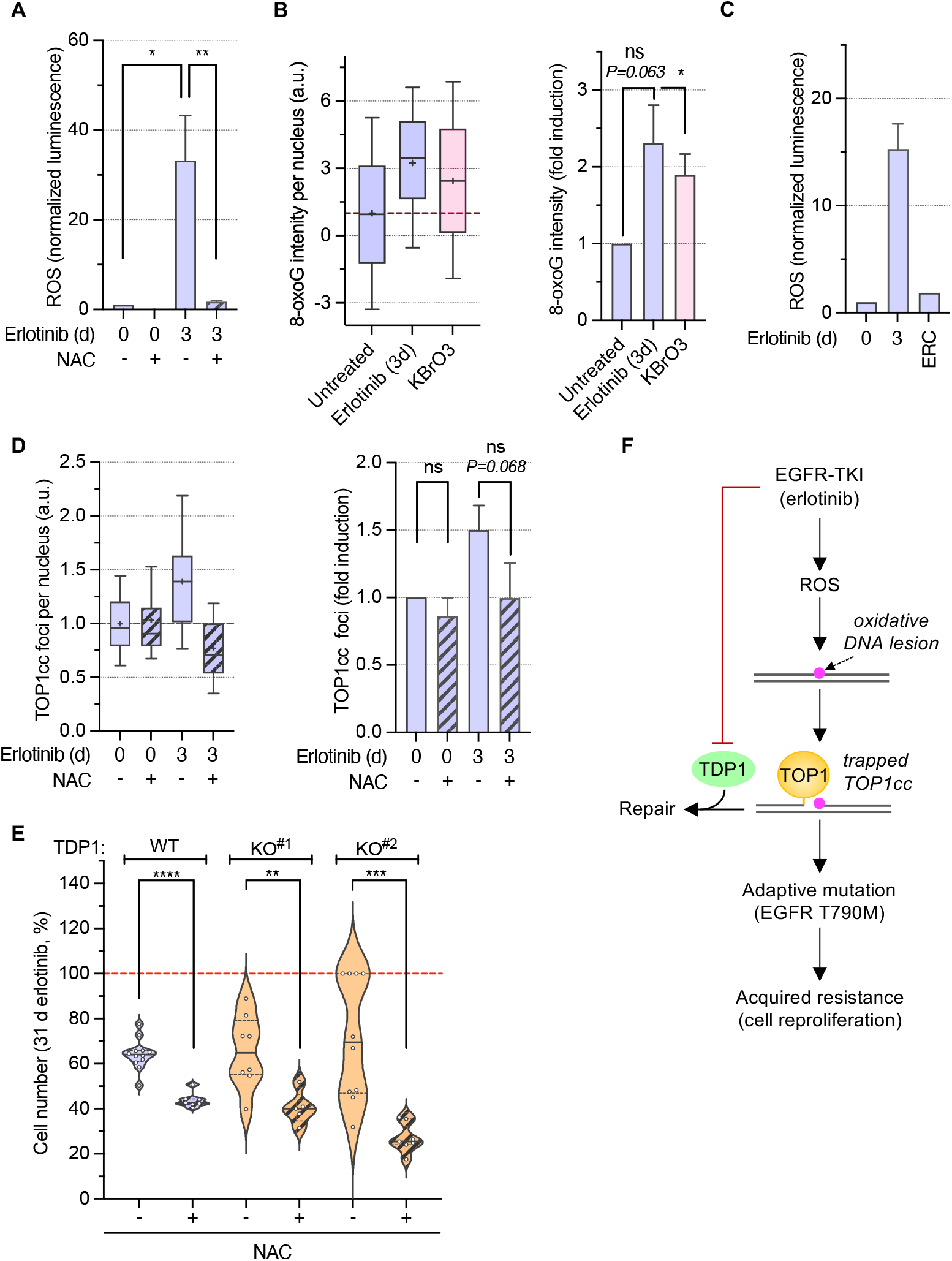
Inhibition of erlotinib-induced ROS prevents TOP1cc accumulation and delays the resumption of cell proliferation. (**A**-**E**) TDP1 WT PC9 cells were treated with erlotinib (1 µM) in the presence or absence of NAC (5 mM) for the indicated times. In panel E, TDP1 KO PC9 cells were also included. (**A**) ROS levels measured using CellROX luminescence, normalized to cell number, and plotted relative to the untreated condition (mean ± SEM; n = 3). \**P* < 0.05, \*\**P* < 0.01 (two-tailed paired t-test on cell number-normalized log2-transformed values). (**B**) 8-oxoG fluorescence intensity per nucleus normalized to untreated cells. Data are shown for a representative experiment (left) and three independent experiments (right: mean ± SEM; n = 3). ns, not significant, \**P* < 0.05 (two-tailed paired t-test on non-normalized log2-transformed values). Treatment with KBrO3 (40 mM, 30 min) was used as a positive control for 8-oxoG induction. (**C**) Same experiment as in (A), showing erlotinib-treated cells (3 days) and DTEC (several weeks of continuous culture with 1 µM erlotinib following relapse) for comparison (mean ± SEM; n = 2). (**D**) Number of TOP1cc foci per nucleus normalized to untreated cells in a representative experiment (left) and in three independent experiments (right: mean ± SEM; n = 3). ns, not significant (two-tailed paired t-test on non-normalized log2-transformed values). (**E**) Violin plot representation of the number of cells after 31 days of erlotinib treatment, monitored by phase-contrast imaging and normalized to untreated cells (100%; dashed red line). Each dot represents one technical replicate (n = 5-12). ns, not significant, \*\**P* < 0.01 \*\*\**P* < 0.001, \*\*\*\**P* < 0.0001 (two-tailed unpaired t-test). (**F**) Model of adaptive resistance induced by EGFR-TKI-mediated TDP1 downregulation.

Altogether, these findings support a model in which erlotinib elevates ROS while transiently downregulating TDP1 in DTCs, thereby promoting TOP1cc accumulation through increased trapping and impaired repair. This ROS-TDP1 axis is consistent with a role for unrepaired TOP1ccs in fostering resistance emergence and subsequent DTC outgrowth (Fig. 4F).

### TDP1 expression is highly heterogeneous and absent in a subset of EGFR-mutated lung cancers

Given our findings implicating TDP1 downregulation in resistance acquisition in EGFR-mutated lung cancer (Figs. 1-4), we next examined TDP1 protein expression across cancer types. Analysis of TDP1 protein expression data from the Cancer Cell Line Encyclopedia (CCLE) revealed substantial heterogeneity across tumor types (Fig. 5A). Notably, NSCLC displayed relatively low TDP1 expression with marked inter-cell line variability (Fig. 5A), consistent with previous reports showing that TDP1 expression can be either lost (*40*) or elevated in lung cancers (*41, 42*).

**Fig. 5.**
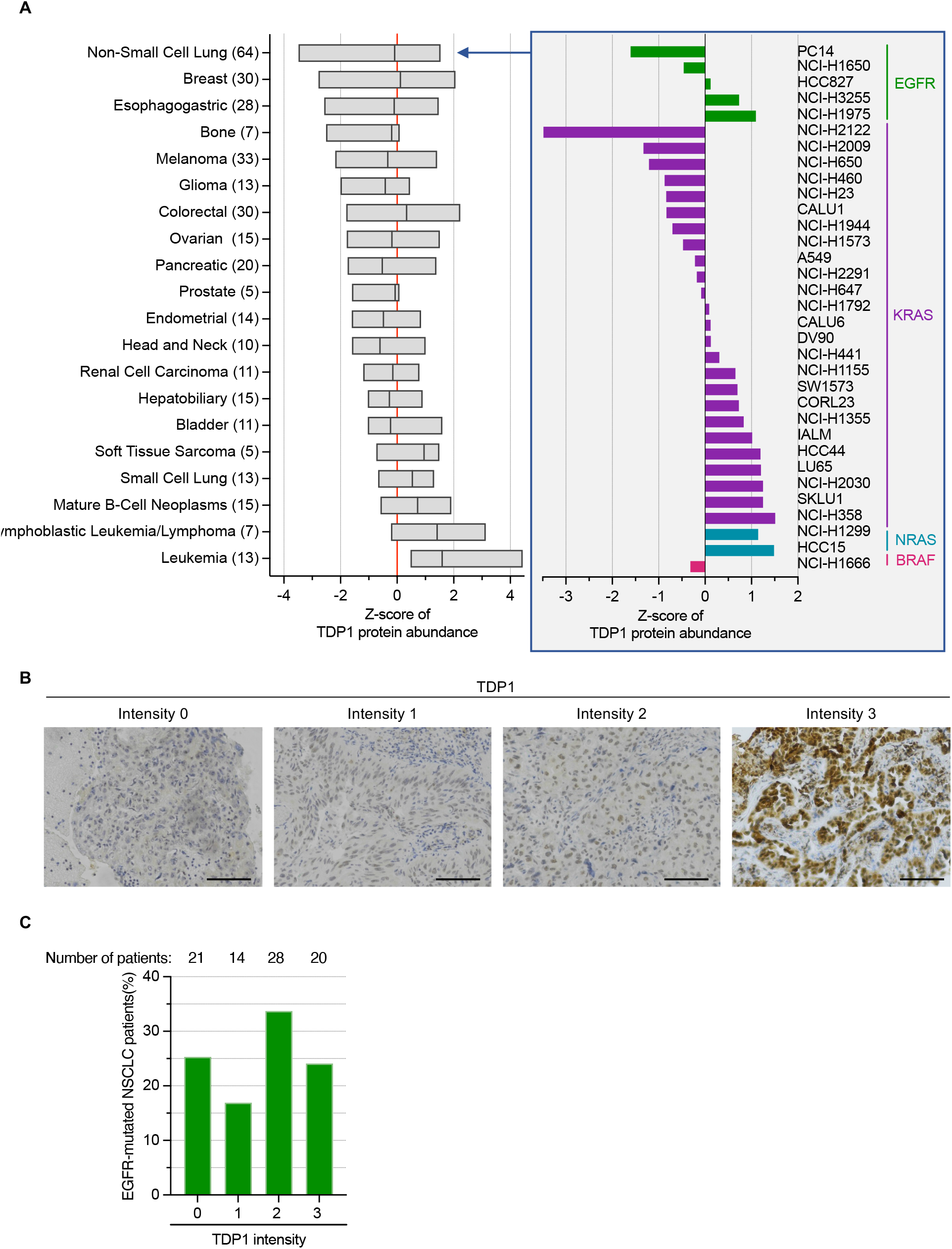
TDP1 protein expression across human tumor cell lines and EGFR-mutated lung cancer biopsies. (**A**) Relative TDP1 protein levels in human tumor cell lines from the Cancer Cell Line Encyclopedia (CCLE; Broad Institute, 2019), grouped by cancer type. Cancer types are sorted from highest to lowest average TDP1 expression. Bars represent the minimum to maximum range, and the line indicates the median. The number of cell lines analyzed per cancer type is indicated in parentheses. The grey box at right highlights the TDP1 z-scores in selected NSCLC cell lines harboring major driver mutations (EGFR, KRAS, NRAS, BRAF). (**B**) Representative immunohistochemistry (IHC) images of TDP1 in 83 biopsies from EGFR-mutated lung cancer patients prior to any treatment. Examples are shown for each expression intensity (0-3), where 0 indicates no detectable TDP1. Scale bars: 100 µM. (**C**) Percentage of biopsies in each intensity category. The number of patient biopsies in each category is indicated.

To determine whether this variability was associated with specific oncogenic drivers, we stratified NSCLC cell lines by EGFR, KRAS, NRAS, or BRAF mutation status. TDP1 expression remained highly heterogeneous within EGFR- and KRAS-mutant groups, for which sufficient numbers of cell lines were available, arguing against a uniform association with a single driver mutation. We next examined the relationship between TDP1 protein and mRNA expression levels. Correlation analysis revealed only a moderate association (r = 0.5) (Fig. S5A), consistent with a previous report in the lung cancer cells from the NCI-60 panel (*40*). This suggests that transcript abundance alone does not fully account for the observed variability in TDP1 protein levels.

To determine whether this heterogeneity extends to primary tumors, we analyzed TDP1 expression by immunohistochemistry (IHC) in tumor biopsies from 83 treatment-naïve patients with EGFR-mutated NSCLC. Notably, 21 of 83 tumors showed no detectable TDP1 signal (Intensity 0; Fig. 5B, C), indicating that approximately one quarter of EGFR-mutated NSCLCs are TDP1-negative by IHC.

### Combined EGFR and TOP1 targeting suppresses resistance acquisition in TDP1-deficient cells

Loss of TDP1 leads to the accumulation of unrepaired TOP1ccs (Fig. 2B,C), which promotes acquisition of the secondary EGFR T790M resistance mutation (Fig. 3A) and confers a proliferative advantage under erlotinib treatment (Fig. 3B-D). We therefore asked whether further increasing TOP1cc levels with topotecan, a specific TOP1cc inducer (*43*), could exceed the cellular tolerance threshold and shift TOP1cc accumulation from a mutagenic to a lethal outcome.

To test this, we first measured TOP1cc levels in PC9 cells exposed to topotecan. Immunofluorescence analysis revealed a concentration-dependent increase in TOP1cc foci (Fig. 6A), consistent with its mechanism of action. We then assessed topotecan sensitivity in TDP1 WT and KO PC9 cells to identify a concentration that had minimal effects on short-term viability in the absence of erlotinib (10 nM; Fig. 6B). Consistent with the IC50 curves, this concentration alone had no detectable effect on cell proliferation in either genotype (Fig. 6C). We next combined this sublethal topotecan concentration with erlotinib and monitor resumption of proliferation. Under combination treatment, reproliferation was observed in TDP1 WT cells but was completely absent in TDP1 KO cells (Fig. 6D, E). These results indicate that further increasing TOP1cc levels with sublethal topotecan shifts the consequences of TOP1cc accumulation in TDP1-deficient cells from mutagenesis to cell death, thereby suppressing resistance outgrowth during erlotinib treatment.

**Fig. 6.**
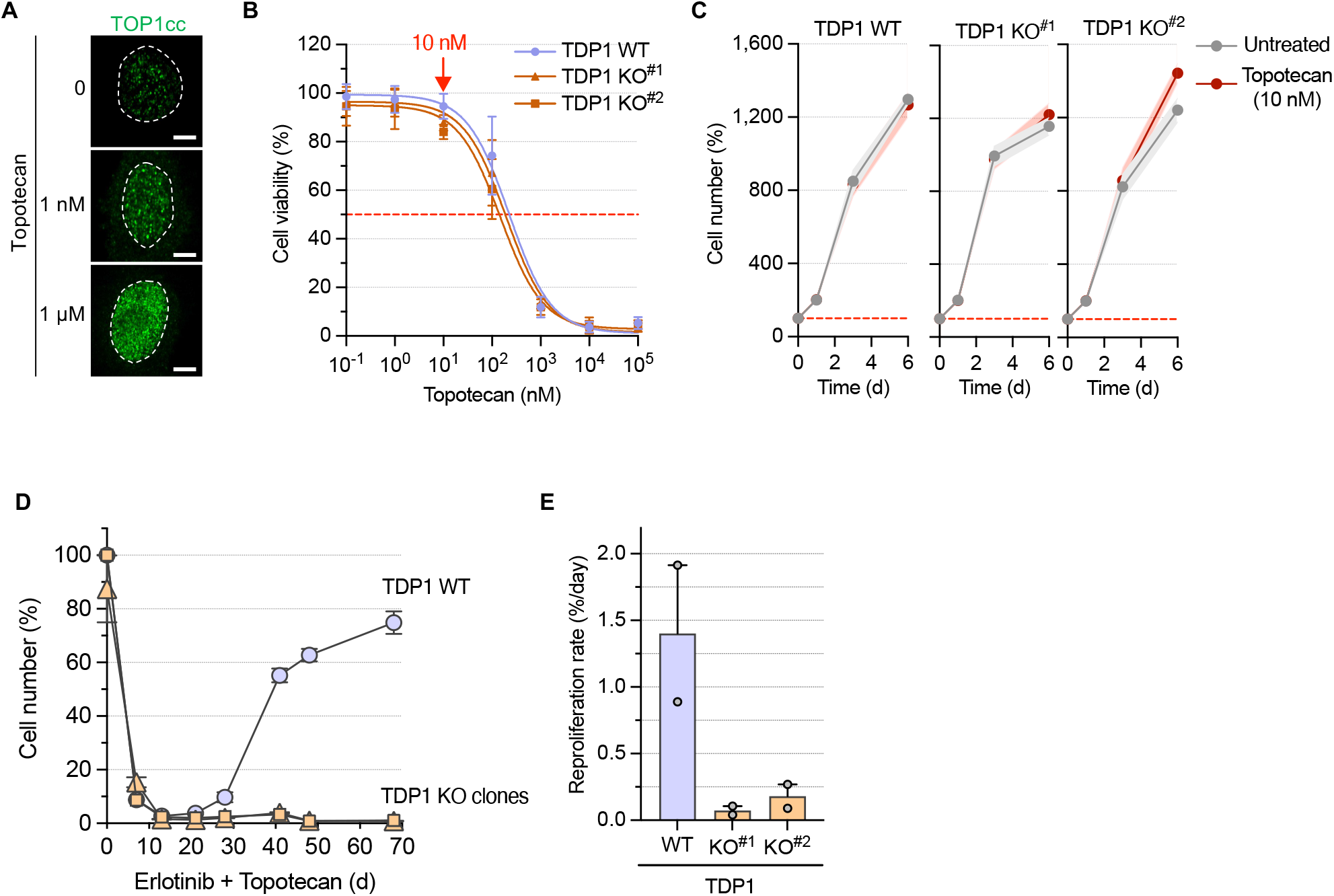
Combination of erlotinib and topotecan prevents re-proliferation of TDP1-deficient cells. (**A**) Representative images of PC9 cells treated with the indicated concentrations of topotecan for 1 h and stained for TOP1ccs and DAPI (DNA). Dashed lines indicate nuclei. Scale bars: 5 µm. (**B**) TDP1 WT and KO PC9 cells were treated with the indicated concentrations of topotecan for 72 h. Cell viability was measured and normalized to untreated cells (100%) (mean ± SEM; n = 4). (**C**) Time-course of TDP1 WT and KO PC9 cell growth, left untreated or treated with 10 nM topotecan. Cell numbers were monitored by phase-contrast imaging and normalized to 100% at the start of the experiment. A representative experiment out of 2-3 independent experiments is shown (mean ± SEM; n = 8 technical replicates). The bold line represents the mean, and the shaded area the SEM. (**D**) Time-course of TDP1 WT and KO PC9 cell number in response to the combination of erlotinib (1 µM) and topotecan (10 nM), monitored by phase-contrast imaging and normalized to untreated cells (100%). A representative experiment out of two independent experiments is shown (mean ± SEM; n = 8 technical replicates). **(E)** Quantification of the reproliferation rate corresponding to the experiments shown in (D) in two independent experiments. The daily reproliferation rate was calculated from the slope of a linear regression performed over the post-nadir phase (days 13 to 38 or 41, depending on the experiment). The nadir was defined as the minimum cell number observed during the DTC stage. Bars represent mean ± SEM from two independent experiments, with individual experimental values shown as dots.

## DISCUSSION

The acquisition of genetic alterations by DTCs is an important mechanism of secondary resistance to targeted therapies (*2-6*). Yet, our understanding of how non-proliferative DTCs acquire such resistance mutations is limited. Here, we identify a transiently mutagenic state in EGFR-TKI-induced DTCs characterized by reduced TDP1-dependent TOP1cc repair and increased oxidative stress, which together promote TOP1cc accumulation and facilitate the acquisition of resistance mutations. Our findings further suggest that this transient defect in TOP1cc resolution may represent a therapeutic vulnerability that could potentially be exploited to limit the emergence and expansion of resistant clones.

The coordinated downregulation of multiple components of the TDP1-mediated TOP1cc repair pathway in DTCs, including TDP1, PARP1, PNKP, XRCC1, and LIG3, argues against selective repression of individual components and instead supports a broader pathway suppression associated with the DTC state. This appears to be linked, at least in part, to cell cycle arrest during DTC entry, as serum deprivation-induced quiescence in WI38-hTERT cells induced a similar, although less pronounced, reduction in the expression of these factors. This interpretation is consistent with previous reports showing reduced expression of several TDP1 pathway components in another model of quiescent cells (*44*) and in senescent cells (*45*). Notably, XRCC1 is a transcriptional target of E2F1 (*44*), a factor downregulated upon cell cycle arrest whose target genes are also repressed in EGFR-TKI-treated DTCs (*9*), providing a potential mechanistic link between cell cycle arrest in DTCs and reduced expression of TOP1cc repair factors. In addition, the more repressive chromatin state reported in DTCs upon EGFR-TKI treatment (*7, 11*) may further contribute to the downregulation of TDP1 pathway components.

Trapped TOP1ccs are a well-established source of genomic instability (*22-24*). In non-proliferative but transcriptionally active cells, persistent TOP1ccs can block the elongating RNA polymerase II (*46-48*), leading to the formation of transcription-dependent DSBs (*30, 49-52*). Consistent with this, defective TOP1cc resolution in TDP1-deficient cells increases the accumulation of transcription-dependent DSBs in non-proliferative cells (*30, 49, 50*). Because these breaks arise in actively transcribed regions (*50*), their inaccurate repair could affect genes whose alteration promotes drug resistance. Although DSBs in transcribed regions are preferentially repaired by homologous recombination (HR) (*53*), this pathway is expected to be impaired in non-replicating DTCs due to both downregulation of HR genes (*13*) and the absence of a sister chromatid as a repair template. Repair may therefore shift toward end-joining pathways, which are generally less accurate than HR. Notably, TDP1 has also been implicated in facilitating non-homologous end joining (NHEJ) (*54-57*), raising the possibility that transient TDP1 downregulation in DTCs may not only increase TOP1cc persistence and the resulting transcription-dependent DSBs, but also impair their repair. This could increase reliance on more error-prone backup pathways such as alternative end-joining (A-EJ), which can operate in G1 (*58*) and when HR and/or NHEJ are compromised (*59-61*). In addition, persistent TOP1ccs can be processed by structure-specific nucleases such as MRE11, CtIP, and XPF, which cleave DNA upstream of the TOP1cc (*25, 62*). Because this pathway generates DNA ends that require gap repair, it may be intrinsically more mutagenic. Together, these observations support a model in which reduced TDP1 activity may increase reliance on alternative TOP1cc processing and error-prone repair pathways in DTCs. Directly testing the contribution of transcription-associated DSB formation, end-joining pathway choice, and nucleolytic TOP1cc removal will be warranted to further define the mechanisms linking impaired TOP1cc resolution to mutagenesis in response to EGFR-targeted therapies.

Our data support the idea that TDP1 downregulation facilitates the acquisition of EGFR T790M during erlotinib treatment. This is consistent with the prominent role of TOP1 at transcriptionally active loci, where it resolves transcription-associated topological stress (*21*), thereby increasing the likelihood of TOP1cc trapping in these regions (*63*), and with T790M being the predominant secondary resistance mutation in response to erlotinib (*12, 34*). Both TDP1 KO clones showed increased T790M emergence relative to WT cells, although the magnitude of this effect differed between clones, with one displaying a more robust and reproducible increase. Notably, the earlier outgrowth of the other clone, despite a more modest increase in T790M suggests that TDP1 loss may not selectively favor this single EGFR mutation, but instead broaden the spectrum of resistance-conferring genetic alterations arising during drug tolerance. In this context, T790M represents a well-characterized and quantifiable readout of mutation acquisition in the PC9/erlotinib model, where resistance is largely driven by recurrent EGFR secondary mutations. By contrast, resistance to later-generation EGFR inhibitors is more heterogeneous and less dominated by recurrent target mutations (*2, 3*), limiting the applicability of comparable single-mutation readouts. It may thus reflect only one component of a broader mutagenic process that could also generate other previously reported resistance alterations, including mutations in genes such as NRAS, RAF1, and BRAF (*12*). Hence, extending these observations to whole-genome analyses in DTCs could reveal the full spectrum of resistance-associated alterations driven by transient TDP1 downregulation. However, previous studies in colorectal and breast cancers have reported that increased mutation burden is not readily detected by whole-exome sequencing in DTCs (*13, 64*), highlighting the intrinsic difficulty of capturing transient, low-frequency, and heterogeneous mutational processes.

Together, our findings support a model in which transient downregulation of the TDP1 pathway, together with increased oxidative stress, promotes TOP1cc accumulation in EGFR-TKI– induced drug-tolerant cells (DTCs), thereby generating a transient mutagenic state that facilitates the acquisition of genetic resistance. Although defined in EGFR-mutated PC9 cells, this mechanism may extend to DTC states induced by other EGFR inhibitors and, more broadly, by diverse targeted therapies and chemotherapies across cancer types, which are frequently associated with metabolic rewiring and oxidative stress (*2, 5, 65*). Cell cycle exit may further contribute to reduced expression of the TDP1-mediated repair pathway (this study and (*44, 45*)) in DTCs, thereby impairing TOP1cc resolution and increasing their accumulation. Our study further identifies impaired TOP1cc resolution as a vulnerability, as shown by the inability of TDP1-deficient DTCs to resume proliferation under combined EGFR inhibition and the FDA-approved TOP1 inhibitor topotecan. This may be particularly relevant in a therapeutic context, as we find that approximately 25% of EGFR-mutated NSCLC lacks detectable TDP1 expression. In this setting, such combination strategies aimed at increasing TOP1cc burden could be further optimized for safety using antibody-drug conjugates (ADCs) delivering topoisomerase I inhibitor payloads, such as SN-38 and exatecan-derived payloads, which are clinically approved (*66*). Further evaluation of this approach will require in vivo validation of combinations between clinically used EGFR-TKIs and topoisomerase I inhibitors, as well as testing across additional targeted therapy settings characterized by low or undetectable TDP1 expression.

## MATERIALS AND METHODS

### Cells and treatments

Human EGFR-mutated non-small cell lung cancer (NSCLC) cell lines were used in this study. PC9 cells (EGFR ΔE746–A750) were provided by Antonio Maraver (IRCM, Montpellier). HCC4006 (CRL-2871; EGFR ΔL747-E749, A750P) were obtained from the American Type Culture Collection (ATCC). To study adaptive resistance, PC9 cells were seeded at 70-80% confluency and treated with 1 µM erlotinib twice a week until relapse. Cells were cultured in RPMI-1640 medium (#509090; Clearline) supplemented with 10% (v/v) fetal bovine serum (FBS) (#S1900-500C; Dutcher) and 1% (v/v) penicillin-streptomycin (#P0781; Sigma) at 37 °C in 5% CO_2_. PC9 and HCC4006 cells were subcloned to minimize the presence of potential pre-existing drug-resistant cells (*9*). WI38 fibroblasts immortalized with hTERT were obtained from Carl Mann (CEA, Gif-sur-Yvette, France). Cells were grown in MEM medium supplemented with 10% (v/v) FBS, 1 mM sodium pyruvate, 2 mM glutamine and 0.1 mM non-essential amino acids at 37 °C in 5% CO_2_. Quiescence was induced by washing the cells twice with serum-free medium and culturing them for 72 h in the growth medium (as above) but with 0.2% (v/v) FBS as previously described (*30, 49, 50*). Cells were routinely tested and confirmed to be free of mycoplasma contamination. Cell numbers were determined using a Countess II automated cell counter (ThermoFisher Scientific). Drugs and chemicals used were erlotinib (#E4997; LC Laboratories), potassium bromate (KBrO3; #309087; Sigma), N-acetyl cysteine (NAC; #A7250; Sigma), ouabain (#O3125; Sigma), and topotecan (#T2705; Sigma). Erlotinib and topotecan were dissolved in DMSO, KBrO3 in phosphate-buffered saline (PBS), and ouabain and NAC in water. Mock samples were treated with the vehicle only.

### Analysis of cell number by real-time high-content microscopy

Cells were seeded in 24- or 48-well plates and maintained under standard culture conditions (37 °C, 5% CO_2_) for the duration of the experiment. For time-lapse analysis, plates were transferred to an Operetta CLS High-Content Imaging System (PerkinElmer) equipped with an environmental control chamber set to 37°C and 5% CO_2_ during image acquisition, using a 20X widefield objective. Plates were imaged immediately after each medium change to exclude detached cells and debris. Cell number per well was quantified with Columbus software (version 2.8.2) or Harmony software (version 4.9). Cell counts were normalized to the first acquisition of each well (day 0), which was set to 100%.

### RNA-seq and data analyses

TDP1 WT PC9 cells were treated with 1 µM erlotinib for 7 days (DTC) and 28 days (DTEC). In parallel, TDP1 KO^#2^ PC9 cells were cultured in the absence of treatment. For each condition, three replicate cultures were generated by splitting cells from a common initial flask and subsequently cultured independently prior to RNA extraction and sequencing. Total RNA was extracted using the RNeasy Plus Mini kit (#74134; Qiagen) according to the manufacturer’s instructions. Library preparation was performed using the TruSeq Stranded mRNA kit (Illumina), and sequencing was carried out on a NovaSeq 6000 with 101 bp paired-end reads at CeGaT GmbH (Tübingen, Germany). Reads were aligned to human genome (GRChg38, Ensembl release 110), and gene expression levels were quantified using RSEM ‘rsem-calculate-expression’ (v1.3.1) with Bowtie2 (v2.5.2). Differential expression analyses were performed using Bioconductor DESeq2 (v1.40.2).

Publicly available RNA-seq data for PC9, HCC827, and NCI-H1975 cells left untreated and treated with 500 nM osimertinib for 21 days (DTC state; GSE193258) (*32*), and for HCC4006 cells left untreated and treated with 1 µM erlotinib for 21 days (DTC state; GSE249721) (*9*) were obtained from the NCBI GEO database. Differential expression analysis was performed using GEO2R.

### Cell extracts and immunoblotting

Cell extracts were obtained by lysing cells for 20 min at 4°C in RIPA buffer (50 mM Tris-HCl pH 8.0, 150 mM NaCl, 1% Triton X-100, 1% sodium deoxycholate, 0.1% SDS, 5 mM EDTA) supplemented with protease and phosphatase inhibitors (Halt Protease & Phosphatase Inhibitor Cocktail; #1861281; ThermoFisher Scientific). Lysates were centrifuged at 16,000 x g for 15 min at 4°C and supernatant was collected. For Figures S2B and S2D, cell extracts were obtained by lysing cells for 15 min at 4°C in buffer containing 10 mM Tris-HCl (pH 7.5) and 1% SDS, supplemented with protease and phosphatase inhibitors (Halt Protease & Phosphatase Inhibitor Cocktail; #1861281; ThermoFisher Scientific). Viscosity of the samples was reduced by sonication. Proteins were separated by SDS-PAGE and immunoblotted with the following antibodies at dilutions recommended by the manufacturer: anti-pan-actin (#MAB1501; Merck-Millipore), anti-DNA ligase 3 (LIG3; #A301-636A; Bethyl), anti-PARP1 (#9542; Cell Signaling Technology), anti-PNKP (#A300-257A; Bethyl), anti-TDP1 (#ab224822; Abcam), anti-TOP1 (#ab109374; Abcam), and anti-XRCC1 (#A300-065A; Bethyl). Membranes were then incubated for 1 h with HRP-conjugated secondary antibodies, either goat anti-mouse (#170-6516; Bio-Rad) or goat anti-rabbit (#170-6515; Bio-Rad). Immunoblots were revealed by chemiluminescence using a ChemiDoc MP Imaging System (Bio-Rad). Protein levels were quantified using ImageLab software (version 6.0.1).

### Cell cycle analysis

Cell cycle analysis was performed as previously described (*30, 67*). Briefly, cells were grown in 96-well plates (CellCarrier Ultra; PerkinElmer) and incubated with 10 µM of the nucleotide analogue EdU for 30 min at 37°C to label newly synthesized DNA. After fixation with 3.7% formaldehyde and permeabilization with 0.5% Triton X-100, incorporated EdU was detected using the Click-iT EdU Alexa Fluor 647 Imaging Kit (#C10340; ThermoFisher Scientific) according to the manufacturer’s protocol. Nuclei were stained with 1 µg/ml Hoechst 33342 provided in the kit for 15 min. Plates were scanned using a 20X objective on an Operetta CLS High-Content Imaging System (PerkinElmer). Cell cycle phases were assigned based on EdU incorporation and Hoechst 33342 intensity using Columbus software (version 2.8.2) or Harmony software (version 4.9): G1 nuclei (EdU-negative, low Hoechst 33342 intensity), S nuclei (EdU-positive), and G2/M nuclei (EdU-negative, high Hoechst 33342 intensity).

### Detection of TOP1ccs by immunofluorescence microscopy

TOP1ccs were detected as previously described (*33*) with modifications (*30*). Briefly, cells were seeded on coverslips in 6-well plates, washed twice with PBS and fixed with 10% formalin for 15 min at 4°C. After three washes with PBS, cells were permeabilized with 0.2% Triton X-100 in PBS for 2 min on ice, followed by an incubation in 0.1% SDS for 5 min at room temperature (RT). Cells were then washed twice in PBS, blocked for 1 h in a buffer containing 10% (w/v) skim milk, 150 mM NaCl and 10 mM Tris-HCl (pH 7.4), and incubated with a mouse anti-TOP1cc antibody (#MABE1084; Merck-Millipore) diluted at 1/1,000 in 5% (v/v) FBS in PBS overnight at 4°C. The next day, cells were washed five times for 4 min each with wash buffer containing 0.1% bovine serum albumin (BSA) and 0.1% Triton X-100 in PBS, and incubated with an anti-mouse antibody coupled to Alexa Fluor 488 (#A-21202; ThermoFisher Scientific) diluted at 1/1,000 in 5% (v/v) FBS in PBS for 1 h at RT. After five washes for 4 min each with wash buffer, coverslips were mounted on glass slides with Duolink *In Situ* Mounting Medium with DAPI (#DUO82040; Sigma) and imaged using an inverted confocal microscope (LSM 780 or 880; ZEISS) with a 63X oil immersion objective. Nuclear foci were analyzed with ImageJ software (version 1.54i).

### Generation of TDP1 knockout PC9 cells

PC9 TDP1 KO cells were generated by targeting exon 5 of TDP1 using CRISPR-Cas9, as previously described (*30*). Cells were co-transfected using jetPEI (#101000053; Polyplus) with i) a plasmid expressing both the eSpCas9 and a gRNA targeting the protospacer on the *ATP1A1* gene (eSpCas9(1.1)_No_FLAG_ATP1A1_G3_Dual_sgRNA plasmid - 86613; Addgene), in which the gRNA targeting *TDP1* gene was inserted (5’-GTTTAACTACTGCTTTGACG-3’); ii) an ATP1A1-RD plasmid donor (86551; Addgene) (*68*), and iii) a single-stranded oligodeoxyribonucleotides (ssODNs) donor for TDP1 (5’-CTT CTT TTC TCC CAT CTA GTT TAA CTA CTG CTT TGA CGT AGA CTG GCT CGT ATA ACA GTA TCC ACC AGA GTT CAG GTG ACG TCC TCA GGG TGA CAG ACA ACA CTA TAA ACT GTA AA-3’) with homology arms for the *TDP1*-targeted site and which contains mutations to insert a stop codon and to mutate the protospacer adjacent motif (PAM) sequence after DSB repair by homologous recombination. Ouabain-resistant cell clones were isolated, screened by immunoblotting and Sanger sequencing. Chromatograms were analyzed using ApE software (https://jorgensen.biology.utah.edu/wayned/ape/) and CRISPR editing was analyzed by TIDE (https://tide.nki.nl).

### Droplet digital PCR for EGFR T790M detection

Droplet digital PCR (ddPCR) was performed to detect the EGFR T790M mutation using primers and fluorescent probes specific for *EGFR* wild-type (HEX) and T790M (FAM) (dHsaMDV2010019; Bio-Rad). Genomic DNA (gDNA) was extracted from cells using the QIAamp UCP DNA Micro kit (#56204; Qiagen) according to the manufacturer’s instructions. Each ddPCR assay included a positive control of gDNA extracted from NCI-H1975 NSCLC cells carrying the *EGFR* T790M mutation, a negative control of gDNA from MCF-7 breast cancer cells, and a water-based handling blank. Samples were run in triplicate in a total reaction volume of 20 µl per well with 8 µl gDNA (50 ng), 10 µl of 2X ddPCR Supermix no dUTP (Bio-rad), 1 µl of the 20X mix of probes and primers (5 µM of each probe and 9 µM of each primer) and 1 µl of sterile water. 20,000 droplets were generated using 20 µl of Droplet Generation Oil in the Droplet Generator (Bio-Rad). Droplets were transferred to a 96-well plate for PCR amplification in a C1000 thermal cycler with the following program: 10 min at 95°C, 40 cycles of 30 sec at 94°C and 1 min at 55°C, followed by 10 min at 98°C. Fluorescence (FAM and HEX) of the droplets was measured using the QX200™ Droplet Reader (Bio-Rad). Only wells with at least 10,000 droplets were included in the analysis. The mean of technical triplicates was calculated using the Poisson distribution. EGFR T790M fractional abundance was calculated as the number of T790M-mutant copies/µl divided by the sum of T790M-mutant and EGFR wild-type copies/µl, multiplied by 100. The limit of detection, calculated by LOB*1.645, was set at 0.066%.

### Detection of reactive oxygen species

Cells were seeded in technical triplicate in white opaque 96-well plates (#6005680; PerkinElmer). H_2_O_2_ levels were quantified using the ROS-Glo H_2_O_2_ Assay (#G8820; Promega) according to manufacturer’s instructions for the non-lytic assay. Briefly, half of the culture medium was transferred to a new white opaque 96-well plate, and the H_2_O_2_ substrate was added to the medium for 6 h before incubation with the detection solution containing luciferase. Luminescence was measured using a CLARIOstar microplate reader (BMG Labtech) and software OPTIMA v1.26. To account for differences in cell number between wells, H_2_O_2_ levels were normalized to cell viability. Cell viability was determined in the original plate (containing the remaining cells) using a luminescence-based ATP assay (ATPlite 1step; #6016731; PerkinElmer) according to the manufacturer’s instructions. Luminescence from the ATP assay was measured as described for the H_2_O_2_ assay. H_2_O_2_ values for each well were divided by the corresponding ATP signal to correct for variations in cell number. Final H_2_O_2_ values are the mean of the three technical replicates.

### Detection of 8-oxoguanine by immunofluorescence microscopy

8-oxoguanine (8-oxoG) was detected as previously described (*69*) with some modifications. Briefly, cells were seeded in 96-well plates (CellCarrier Ultra; PerkinElmer) and treated with erlotinib. As a positive control for 8-oxoG induction, cells were treated with 40 mM KBrO3 for 40 min. After treatment, cells were washed three times with PBS, fixed in acetone/methanol (1/1) for 4 min at 4°C, and air-dried. Cells were then rehydrated in PBS for 15 min, and DNA was denatured with 2 N HCl for 45 min at RT. After three washes in PBS, HCl was neutralized for 5 min with 50 mM Tris-HCl pH 8.8 in PBS at RT, and cells were permeabilized with 0.1% Triton X-100 in PBS for 5 min. Cells were then incubated in blocking buffer (0.1% Triton, 3% BSA, 1% normal goat serum (G9023; Sigma) in PBS) for 1 h at 37°C, and incubated with a mouse anti-8-oxoG antibody (ab48508; Abcam) diluted a 1/2000 in blocking solution overnight at 4°C. The next day, cells were washed three times (10 min each) with 0.1% Triton X-100 in PBS and incubated with an anti-mouse antibody coupled to Alexa Fluor 488 (#A-21202; ThermoFisher Scientific) diluted at 1/1,000 in 0.1% Triton X-100 in PBS for 1 h at 37°C. After three washes (10 min each) with 0.1% Triton X-100 in PBS, nuclei were stained with 1 µg/ml Hoechst 33342 (ThermoFisher Scientific) for 15 min, washed twice with PBS and stored at 4°C until imaging. Plates were imaged with a 20X objective using an Operetta CLS High-Content Imaging System (PerkinElmer) with Harmony software (version 4.9), and imaged were analyzed with Harmony software (version 4.9) as described (*67*). Nuclei were segmented based on Hoechst staining, and total fluorescence intensity was extracted for each nucleus from the 8-oxoG channel. Background signal, estimated from secondary antibody-only controls, was subtracted from all measurements, and the resulting values were normalized to nuclear area. To compare independent experiments, the 8-oxoG intensity was normalized in each individual experiment as indicated in the figure legends.

### CCLE data base analysis

Publicly available data from the Cancer Cell Line Encyclopedia (CCLE; Broad Institute, 2019), were used to evaluate TDP1 protein expression across cancer cell lines. Protein abundance was obtained from CCLE mass spectrometry-based proteomics and reported as Z-scores relative to bridge samples. Within NSCLC subset, cell lines were stratified according to major oncogenic driver alterations, based on annotations provided in DepMap (https://depmap.org/portal/cell_line/). To assess concordance between transcript and protein expression, Z-scores of TDP1 mRNA expression relative to all samples (log RNA Seq RPKM) was obtained from CCLE 2019. Pearson’s correlation was assessed on 64 NSCLC samples with both TDP1 protein and mRNA level data. Proteomics and transcriptomics datasets were downloaded from the DepMap portal (https://depmap.org/portal).

### Tissue samples and TDP1 immunohistochemistry

Tissue samples correspond to the remaining parts of whole tumor tissues from patients, obtained from the tumor library of the CHU Biological Resource Center (IUCT-O), declared to the Ministry of Research under number DC-2008-463. All clinical, pathological, and molecular data were prospectively collected. The study was performed using formalin-fixed, paraffin-embedded (FFPE) tumor tissues from 83 patients diagnosed with NSCLC. All patients harbored EGFR mutations (exon 19 or exon 21). These tissues were collected from surgical resections performed prior to EGFR-TKI treatment and were used for TDP1 immunohistochemistry (IHC) analyses.

TDP1 IHC was performed as previously described (*70*) on 3-µm FFPE NSCLC tissue sections derived from these patients. Epitope retrieval was performed using Flex Target Retrieval high pH Solution for 20 min in a PT Link system (Agilent). Endogenous peroxidase activity was quenched, and sections were incubated with anti-TDP1 antibody (#Ab224822; Abcam) for 1 h at 1/200 dilution. Detection was performed using the EnVision system on an Autostainer (Agilent) according to the manufacturer’s instructions. Hematoxylin was used as counterstain. Antibody specificity was validated using TDP1 WT and TDP1 KO PC9 cells as controls. Two operators, blinded to clinical data, independently evaluated IHC staining scores.

### Cell viability assays

Cells were seeded in technical triplicate in 96-well plates and treated for 3 days. Cell viability was assessed using CellTiter 96® AQueous One Solution Cell Proliferation Assay (MTS; #G3580; Promega) according to manufacturer’s instructions. Absorbance at 490 nm was measured using a CLARIOstar microplate reader (BGM Labtech) with OPTIMA software (v1.26). Absorbance values were corrected by subtracting background signal measured in wells containing reagent only. Corrected absorbance values were normalized to untreated cells, which were set to 100%. Final values are the mean of the three technical replicates.

### Statistical Analysis

Information on biological replicates (n) is provided in the figure legends. Statistical analyses were performed using GraphPad Prism software (versions 9-11). “Ns” indicates no significant difference. A *P* value < 0.05 was considered statistically significant. \**P* < 0.05, **P < 0.01, \*\*\**P* < 0.001, \*\*\*\**P* < 0.0001. The exact *P* value is indicated when it is close to 0.05 but not significant.

## ACKNOWLEDGMENTS

We thank Antonio Maraver for providing PC9 cells, Carl Mann for WI38-hTERT cells, and Nicolas Bery for providing cloning tools and advice. We also thank Anna Campalans for sharing the protocol for 8-oxoG detection by immunofluorescence microscopy. We are grateful to Lara Fernandez Martinez for assistance with RNA extraction, Aurélia Doussine and Sarah Figarol for technical assistance and analysis of ddPCR experiments, respectively, and Latré Lawson Body for technical assistance. Finally, we acknowledge the CRCT platform and Laetitia Ligat for support with microscopy.

## Author contributions

Conceptualization: MG, OS. Investigation: MG, RG, ACa, ETC, SS, NO, CD, EB, DP, CT, ACr. Methodology: MG, ETC, SS, NO, CD, JM, TF, OC, GF, AP, OS. Data curation: MG, RG, ECT, SS, NO, MV, CD, EB, AL, MM, OC, AP, OS. Validation: MG, OS. Project administration: MG, OS. Supervision: OS. Formal analysis: MG, ECT, NO, MV, AL, MM, AP, OS. Writing-original draft: MG, OS. Writing-review & editing: MG, RG, ETC, DP, ACr, JM, OC, GF, AP, OS.

## Data and materials availability

RNA-seq datasets generated in this study are available at the Gene Expression Omnibus under accession code GSE330247. The datasets include PC9 cells untreated and treated with 1 µM erlotinib for 7 days (DTC) and 28 days (DTEC), as well as PC9 TDP1 KO^#2^ cells. PC9 TDP1 KO cells (KO^#1^ and KO^#2^) generated in this study are available without restrictions from the corresponding author upon request. All data are available in the main text or the supplementary materials.

