## Supplemental Figures S1 to S5 for "Persistent TOP1 cleavage complexes in drug-tolerant cells drive adaptive resistance to EGFR-targeted therapies in lung cancer"

Mathéa Geraud *et al.*

**This PDF file includes:** Figs. S1 to S5

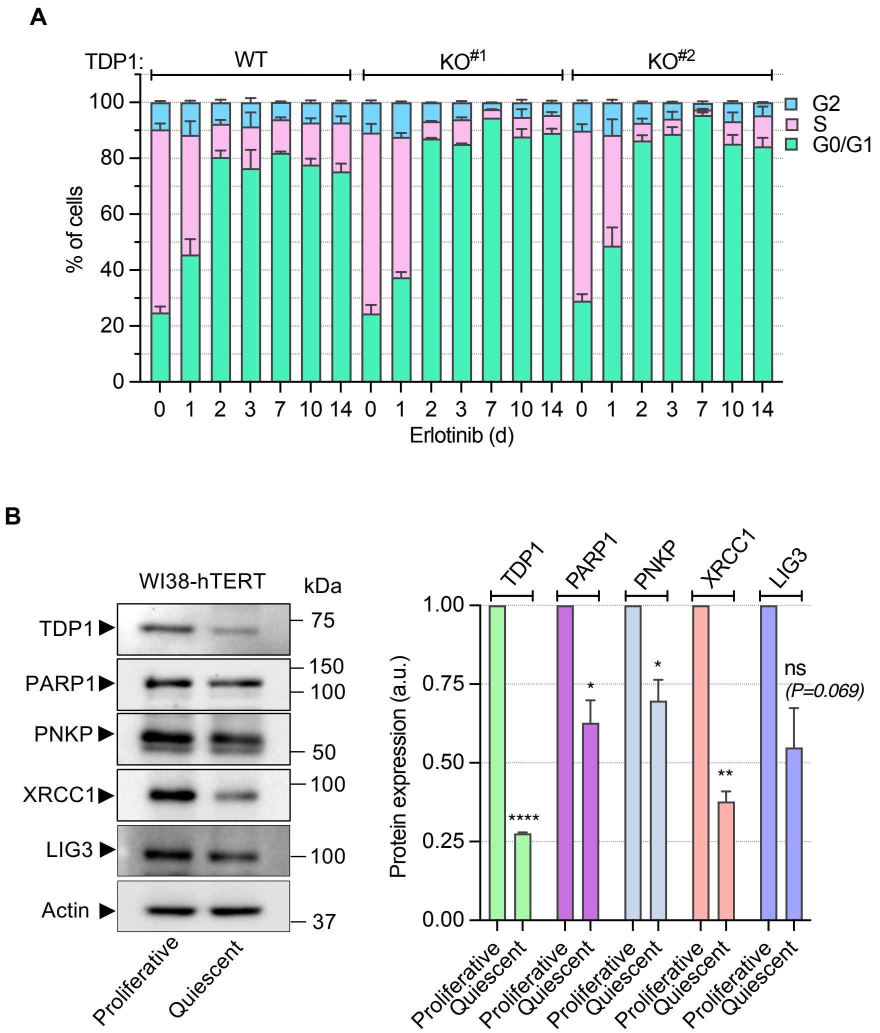


**Fig. S1. Cell cycle analyses in PC9 cells upon erlotinib treatment and expression of TDP1 complex components in proliferative vs quiescent cells.**
(**A**) TDP1 WT and KO PC9 cells were treated with erlotinib (1 µM) for the indicated times. At the end of the treatment, cells were incubated with EdU (10 µM) for 30 min before staining for EdU and Hoechst 33342 (DNA). The percentages of cells in the cell cycle phases were identified by Hoechst 33342 intensity and EdU incorporation; G1 (EdU-negative and low Hoechst 33342), G2 (EdU-negative and high Hoechst 33342) (mean ± SEM; n = 2-8). (**B**) Western blot of proteins of the TDP1 complex in proliferative and quiescent (serum-starved) WI38-hTERT cells. Actin: loading control. Left: representative blots. Right: quantification normalized to actin (mean ± SEM; n = 3). ns, not significant, **P* < 0.05, ***P* < 0.01, *****P* < 0.0001 (one-sample t-test vs. 1).

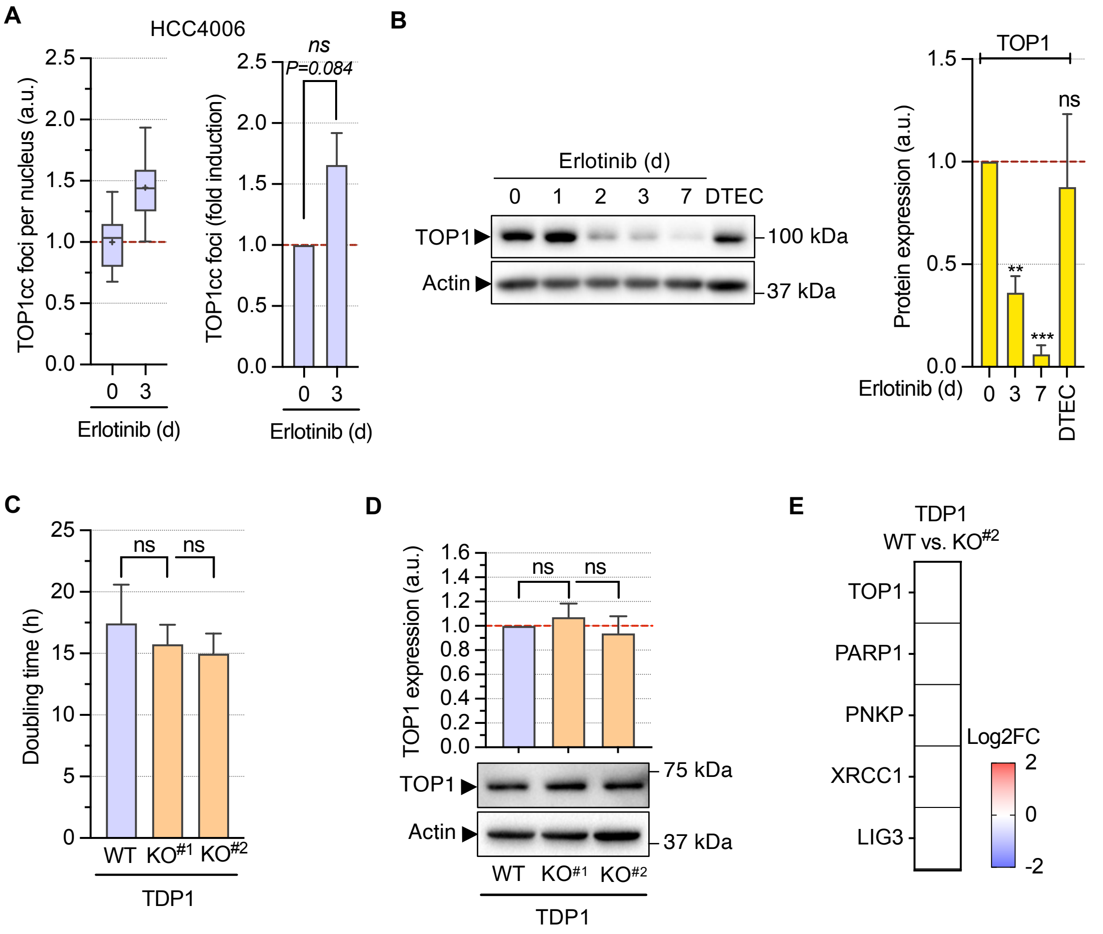


**Fig. S2. TDP1 loss does not alter TOP1 expression or proliferation in lung cancer cells.** (**A**) HCC4006 cells were treated with erlotinib (1 µM) for 3d and stained for TOP1ccs and DAPI (DNA). The number of TOP1cc foci per nucleus normalized to untreated cells is shown in a representative experiment (left) and in three independent experiments (right: mean ± SEM; n = 3). ns, not significant, (two-tailed paired t-test on non-normalized log2-transformed values). (**B**) Western blot of TOP1 in PC9 cells treated with erlotinib (1 µM) for the indicated times. DTEC corresponds to several weeks of continuous culture with erlotinib (1 µM) following relapse. Actin: loading control (which is from the same experiment as in Fig. 1E). Left: representative blot. Right: quantification normalized to actin (mean ± SEM; n = 3). ns, not significant, $^{**}P < 0.01$, $^{***}P < 0.001$ (one-sample t-test vs. 1). (**C**) TDP1 WT and KO PC9 cells were monitored overtime by phase-contrast imaging and doubling time was calculated from the exponential phase of the growth curves (mean ± SEM; n = 3). ns, not significant (two-tailed paired t-test). (**D**) Western blot of TOP1 in TDP1 WT and KO PC9 cells. Actin: loading control. Bottom: representative blot. Top: quantification normalized to actin (mean ± SEM; n = 3). ns, not significant. (**E**) Differential expression of TOP1 and other TDP1 complex genes (TDP1 itself is not shown) in TDP1 KO#2 versus WT PC9 cells. Heatmap shows log2 fold changes (log2FC) of transcript levels, with white boxes indicating non-significant changes (adjusted $P$-value ≥ 0.05).

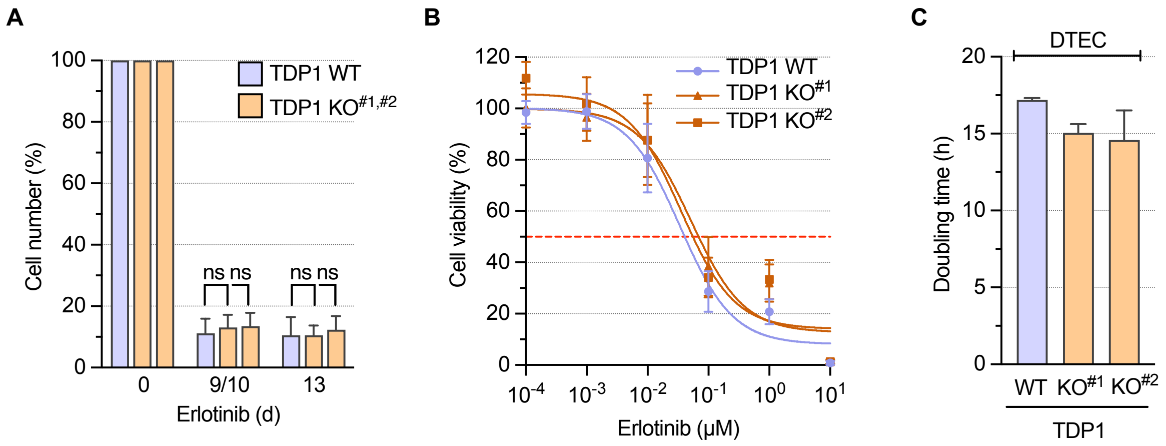


**Fig. S3. TDP1 KO does not alter PC9 cell growth, erlotinib response, or doubling time of erlotinib-resistant cells.**

(**A**) TDP1 WT and KO PC9 cells were treated with erlotinib (1 μM) and cultured over time. Cell numbers were counted on days 9 or10 and 13 (DTC state) by phase-contrast imaging, and normalized to untreated cells (100%) (mean ± SEM; n = 3). ns, not significant (two-tailed paired t-test). (**B**) TDP1 WT and KO PC9 cells were treated with the indicated concentrations of erlotinib for 72 h. Cell viability was measured and normalized to untreated cells (100%) (mean ± SEM; n = 3). (**C**) Erlotinib-resistant TDP1 WT and KO PC9 cells (DTEC) were monitored over time by phase-contrast imaging and doubling time was calculated from the exponential phase of the growth curves (mean ± SEM; n = 2).

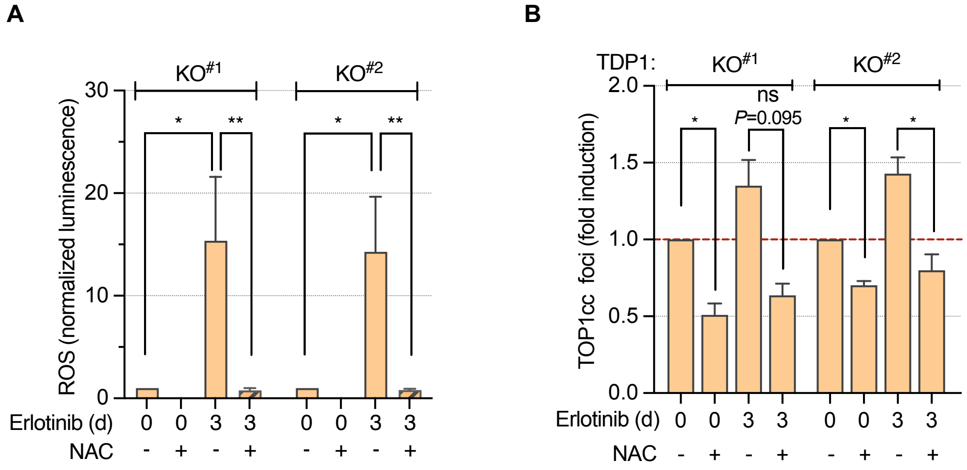


**Fig. S4. Erlotinib induces ROS and ROS-dependent TOP1ccs in PC9 TDP1 KO cells.**
(**A,B**) PC9 TDP1 KO cells were treated with erlotinib (1 μM) in the presence or absence of NAC (5 mM) for 3d. (**A**) ROS levels were measured using CellROX luminescence, normalized to cell number, and plotted relative to the untreated condition (mean ± SEM; n = 3). **P* < 0.05, ***P* < 0.01 (two-tailed paired t-test on cell number-normalized log2-transformed values). (**B**) Number of TOP1cc foci per nucleus normalized to untreated cells (mean ± SEM; n = 3). ns, not significant, **P* < 0.05 (two-tailed paired t-test on non-normalized log2-transformed values).

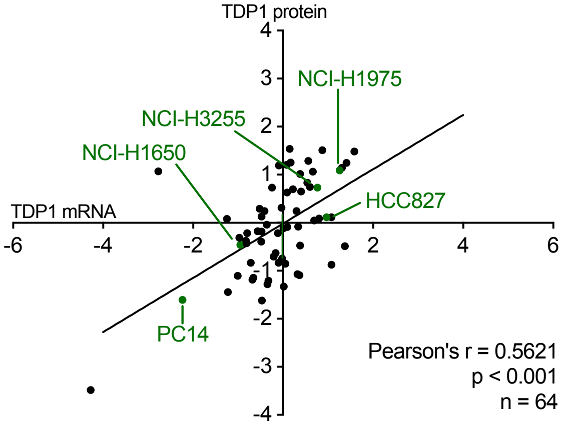


**Fig. S5. Correlation between TDP1 protein and mRNA levels in NSCLC cell lines from the CCLE.**

The 64 NSCLC cell lines are shown, with TDP1 protein and mRNA expression levels displayed as z-scores calculated across all CCLE cell lines. Green dots indicate EGFR-mutated NSCLC cell lines.
